# Rapid fall in circulating non-classical monocytes in ST elevation myocardial infarction patients correlates with cardiac injury

**DOI:** 10.1101/2021.02.01.428825

**Authors:** Sarah A. Marsh, Catherine Park, Rachael E. Redgrave, Esha Singh, Lilia Draganova, Stephen E. Boag, Luke Spray, Simi Ali, Ioakim Spyridopoulos, Helen M. Arthur

## Abstract

**Objective:** Myocardial infarction leads to a rapid innate immune response that is ultimately required for repair of damaged heart tissue. We therefore examined circulating monocyte dynamics immediately after reperfusion of the culprit coronary vessel in STEMI patients to determine whether this correlated with level of cardiac injury. A mouse model of cardiac ischaemia/reperfusion injury was subsequently used to establish the degree of monocyte margination to the coronary vasculature that could potentially contribute to the drop in circulating monocytes.

**Approach and Results:** We retrospectively analysed blood samples from 51 STEMI patients to assess the number of non-classical (NC), classical and intermediate monocytes immediately following primary percutaneous coronary intervention. Classical and intermediate monocytes showed minimal change. On the other hand circulating numbers of NC monocytes fell by approximately 50% at 90 minutes post-reperfusion. This rapid decrease in NC monocytes was greatest in patients with the largest infarct size (p<0.05) and correlated inversely with left ventricular function (r=0.41, p=0.04). The early fall in NC monocytes post reperfusion was confirmed in a second prospective study of 13 STEMI patients. Furthermore, in a mouse cardiac ischaemia model, there was significant monocyte adhesion to coronary vessel endothelium at 2 hours post-reperfusion pointing to a specific and rapid vessel margination response to cardiac injury.

**Conclusions:** Rapid depletion of NC monocytes from the circulation in STEMI patients following coronary artery reperfusion correlates with the level of acute cardiac injury and involves rapid margination to the coronary vasculature.

**Graphical Abstract:** 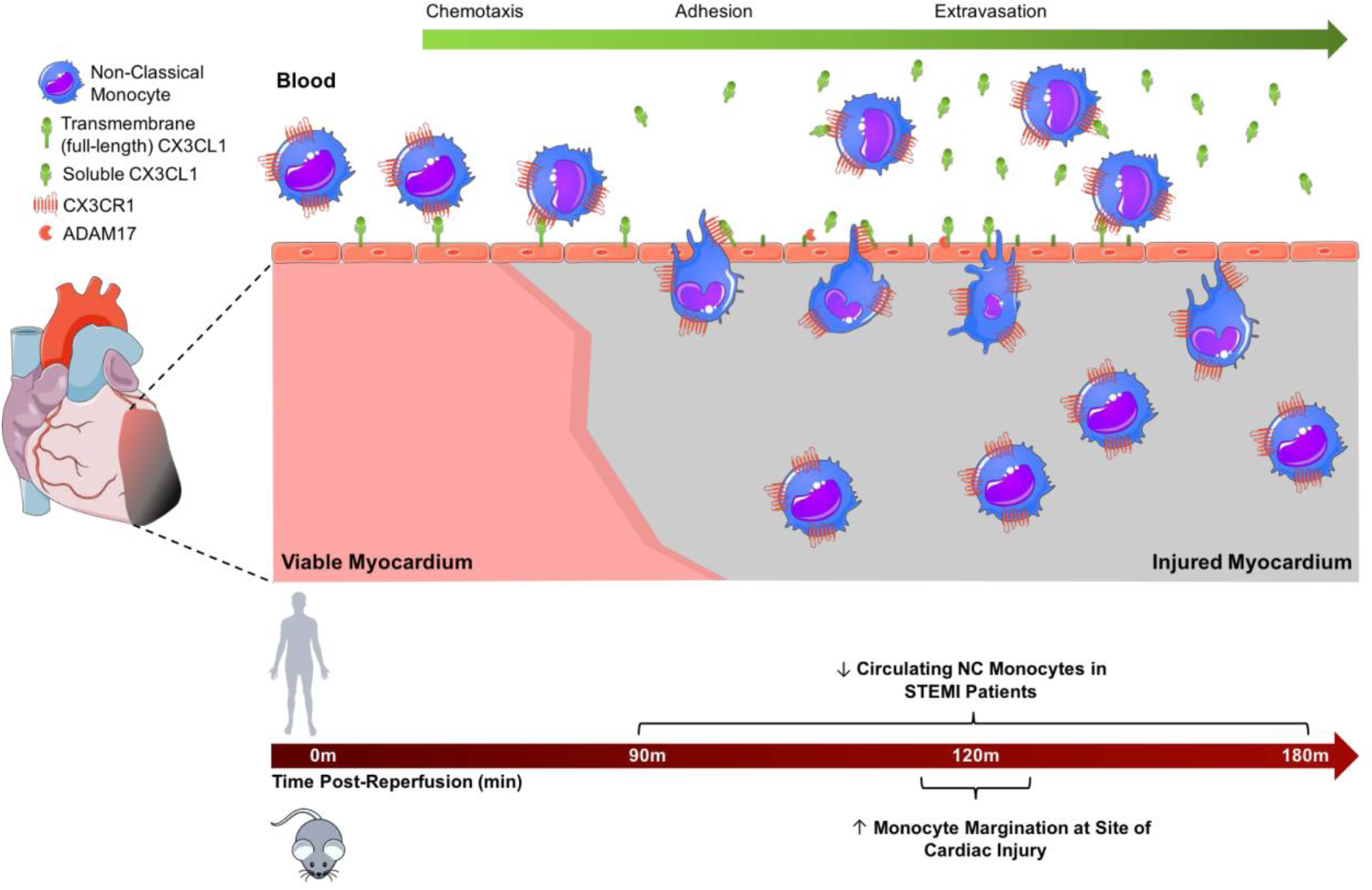

**Highlights:**

- **3-5 bullet points that summarize the major findings of the study.**

1. Circulating non classical monocytes show a rapid fall in STEMI patients within 90 minutes of re-opening the culprit coronary artery.
2. The extent of the drop in non classical monocytes correlates with loss of cardiac function and increased infarct size.
3. A mouse model of cardiac ischaemia and reperfusion shows rapid margination of monocytes to the coronary vasculature

## Introduction

Acute myocardial infarction (MI) is a leading cause of mortality. The advent of primary percutaneous coronary intervention (PPCI) has improved survival rates following MI, but patients are at increased risk of developing subsequent major adverse cardiac events (MACE)^1^. Finding interventions to reduce this risk depends on successfully addressing three inter-dependent goals: to better understand the detrimental processes that contribute to increased MACE risk, to therapeutically intervene to attenuate these events, and to identify robust biomarkers for those patients at highest risk so they can be prioritised for therapeutic interventions.

Biomarkers that accurately reflect the degree of myocardial damage are critical for patient stratification. Conventionally this corresponds to cardiac troponin released from injured cardiomyocytes, but this does not reach a peak until up to 12 hours after initiation of cardiac injury. In contrast, the acute inflammatory response is triggered within minutes of cardiac tissue damage. Ischaemic cardiomyocytes release reactive oxygen species (ROS) upon re-oxygenation after reperfusion and dying cardiomyocytes release damage-associated molecular patterns (DAMPs). Both ROS and DAMPs trigger recruitment of leukocytes to the injured heart to initiate the repair process ^2^. In light of the rapidity of these responses there is considerable interest in using dynamic changes in circulating leukocytes as biomarkers to predict the risk of subsequent MACE. For example, we have previously shown that the percentage fall in circulating effector T cells, during the first 90 minutes after reperfusion of ST elevation STEMI patients is significantly correlated with the degree of microvascular occlusion (MVO) that impacts patient outcomes in the longer term^3^. The cytokine Fractalkine (also known as CX3CL1) was identified as a key mediator in this process. As monocytes are critical components of the repair process, and their properties vary in accordance with expression of the fractalkine receptor (CX3CR1) we asked whether circulating numbers of these different monocyte populations also show dynamic changes in STEMI patients immediately following reperfusion and whether these correlated with patient outcomes.

Human monocyte subpopulations have been defined as classical (CD14^++^ CD16^-^), non-classical/NC (CD14^+^ CD16^++^) and intermediate (CD14^++^ CD16^+^)^4^. Classical monocytes also express CCR2 and are associated with inflammation, whilst NC monocytes express high levels of CX3CR1, and their murine equivalent has been shown to have an endothelial surveillance role^5, 6^. Previous studies have revealed an association between monocyte dynamics at one or more days after PPCI and patient outcomes. Higher numbers of circulating classical monocytes in STEMI patients at 24h following hospital admission are found in patients who progress to develop adverse cardiovascular events by two years^7^, whilst increased numbers of circulating classical monocytes in the first few days post PPCI are associated with increased infarct size^8,9^ and reduced myocardial salvage^10^. In addition, increased numbers of circulating intermediate monocytes from 24h post-reperfusion correlate with increased troponin levels^11^, and positively associate with two-year MACE^7^. Thus, increased numbers of classical and intermediate monocytes in the first few days after PPCI are associated with an increased risk of poorer patient outcomes. However, by the time these blood measurements are being made there has been additional release of monocytes from splenic and bone marrow stores, secondary to the initial acute intra-cardiac immune response ^12, 13^. Thus, any primary changes in the circulating monocyte numbers that are directly driven by the initial cardiac injury will be masked. To our knowledge no studies to date have focused on the dynamics of circulating monocytes at early time points (minutes to hours) after reperfusion, which would more closely reflect the primary response of monocytes to cardiac ischaemia and reperfusion. We therefore asked whether the dynamics of circulating monocytes in acute STEMI patients at the time of, and immediately following, PPCI could provide useful biomarkers of cardiac outcomes. Subsequently, we used a surgical mouse model of cardiac ischemia/reperfusion to track changes in monocyte location within the coronary vessels after reperfusion.

## Materials and Methods

Ethical approval for the patient studies was obtained from the National Research Ethics Service Committee North East (REC references: 12/NE/0322 and 16/NE/0405). Both studies were conducted according to the principles set out in the Declaration of Helsinki and written informed consent was obtained from all patients.

### Retrospective Analysis of STEMI patient Flow Cytometry Data

Analysis of previously obtained flow cytometry data^3^ was used to quantify circulating monocyte subpopulations in 51 STEMI patients. Blood samples were taken at the time of PPCI and at 15 minutes, 30 minutes (culprit coronary arterial blood), 90 minutes (radial artery) prior to removal of the angioplasty sheath, and at 24 hours (venous blood) post-reperfusion. A further venous blood sample was collected between 3 and 6 months later in 20 of these patients. Inclusion criteria for STEMI patients were chest pain of onset within 6 hours and new ST segment elevation. Exclusion criteria included cardiogenic shock, previous MI, active infection or malignancy, chronic inflammatory conditions and patent arterial flow in the infarct related artery. A total of 42 patients underwent cardiac MRI scanning at 1–8 days after infarction as previously described^3^ to determine infarct size, cardiac function and MVO. A group of 15 NSTEMI patients undergoing non-emergency PPCI was also recruited for comparison (Table 1). Immunolabelled blood leukocytes were analyzed using a BD FACSCanto II and FlowJo acquisition software. Antibody details are listed in Supplemental Materials. The absolute cell counts of each monocyte subset were calculated in conjunction with absolute counts for the parent monocyte population in a TruCount assay (Supplementary Figure I).

**Table 1.**
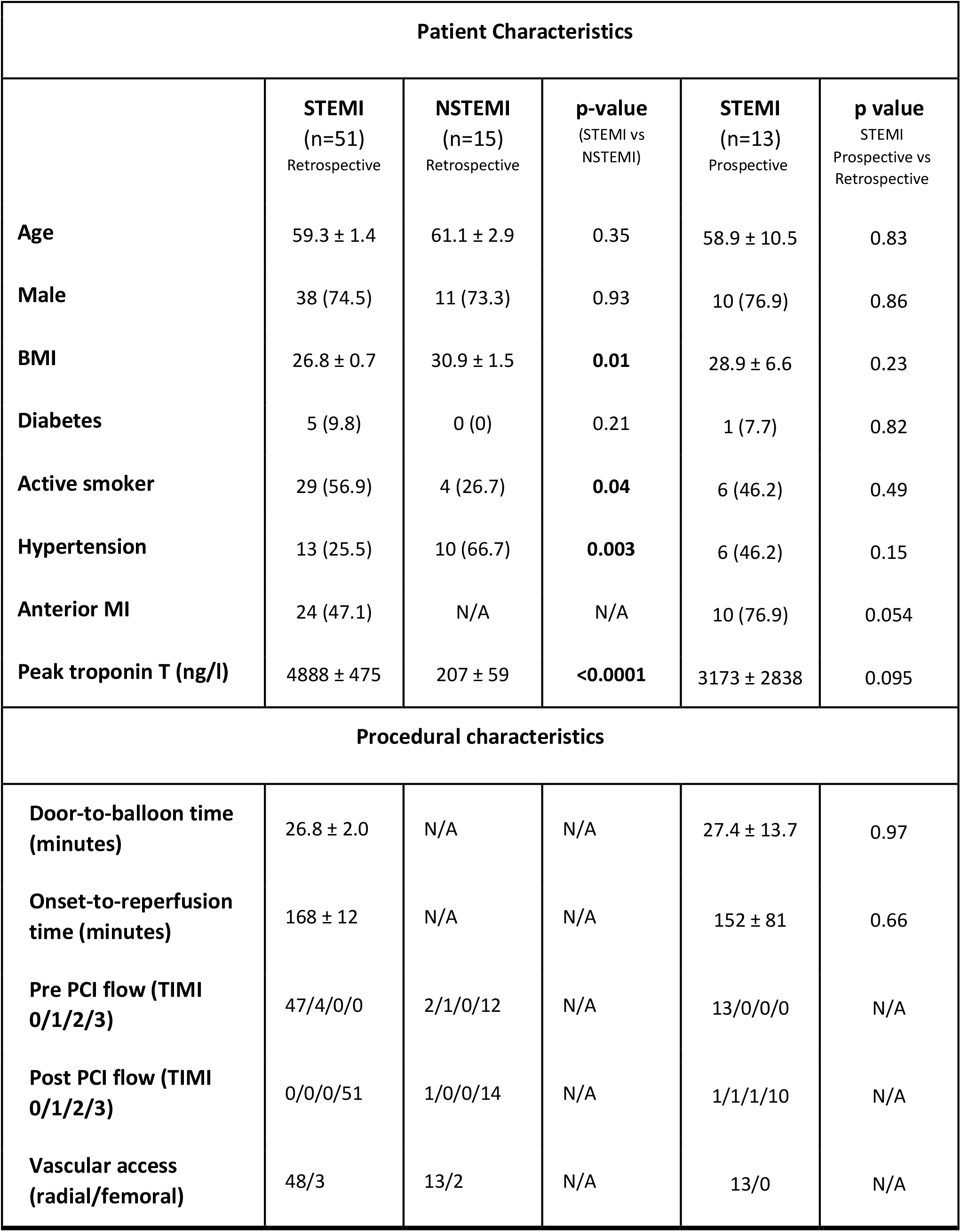
Comparison of baseline data for STEMI and NSTEMI patients in the retrospective study and STEMI patients in the prospective study. Continuous variables expressed as mean ± SEM; categorical variables expressed as n (%). Statistical analyses were performed using unpaired, two-tailed t-tests for continuous data, and Chi-squared tests for categorical data.

| Patient Characteristics |  |  |  |  |  |
| --- | --- | --- | --- | --- | --- |
|  | STEMI<br>(n=51)<br>Retrospective | NSTEMI<br>(n=15)<br>Retrospective | p-value<br>(STEMI vs<br>NSTEMI) | STEMI<br>(n=13)<br>Prospective | p value<br>STEMI<br>Prospective vs<br>Retrospective |
| Age | 59.3 ± 1.4 | 61.1 ± 2.9 | 0.35 | 58.9 ± 10.5 | 0.83 |
| Male | 38 (74.5) | 11 (73.3) | 0.93 | 10 (76.9) | 0.86 |
| BMI | 26.8 ± 0.7 | 30.9 ± 1.5 | **0.01** | 28.9 ± 6.6 | 0.23 |
| Diabetes | 5 (9.8) | 0 (0) | 0.21 | 1 (7.7) | 0.82 |
| Active smoker | 29 (56.9) | 4 (26.7) | **0.04** | 6 (46.2) | 0.49 |
| Hypertension | 13 (25.5) | 10 (66.7) | **0.003** | 6 (46.2) | 0.15 |
| Anterior MI | 24 (47.1) | N/A | N/A | 10 (76.9) | 0.054 |
| Peak troponin T (ng/l) | 4888 ± 475 | 207 ± 59 | **<0.0001** | 3173 ± 2838 | 0.095 |
| Procedural characteristics |  |  |  |  |  |
| Door-to-balloon time<br>(minutes) | 26.8 ± 2.0 | N/A | N/A | 27.4 ± 13.7 | 0.97 |
| Onset-to-reperfusion<br>time (minutes) | 168 ± 12 | N/A | N/A | 152 ± 81 | 0.66 |
| Pre PCI flow (TIMI<br>0/1/2/3) | 47/4/0/0 | 2/1/0/12 | N/A | 13/0/0/0 | N/A |
| Post PCI flow (TIMI<br>0/1/2/3) | 0/0/0/51 | 1/0/0/14 | N/A | 1/1/1/10 | N/A |
| Vascular access<br>(radial/femoral) | 48/3 | 13/2 | N/A | 13/0 | N/A |

### Prospective STEMI Patients and Blood Sampling

A fresh cohort of 13 STEMI patients undergoing PPCI at the Freeman Hospital, Newcastle-upon Tyne were recruited to the prospective study (Table 1). Inclusion criteria were chest pain of onset within 6 hours and new ST segment elevation. Exclusion criteria included cardiogenic shock, previous MI, active infection or malignancy, chronic inflammatory conditions or patent arterial flow in the infarct related artery. Coronary angiography and PPCI were performed, and arterial blood was acquired at the start of the procedure from the aorta and culprit artery, then at 90 and 180 minutes following reperfusion from the radial artery and venous blood was obtained at 24 hours.

### Flow cytometry of bloods from prospective STEMI patients

Blood was obtained in 4 ml EDTA tubes (BD Biosciences). Absolute counts of total monocytes in whole blood were obtained using trucount beads (BD Biosciences, #340334). Monocyte subpopulations were determined in parallel using an eight-color flow cytometric assay. At each time point 50μl of whole blood was stained with fluorescently conjugated antibodies to detect CD14, CD16, HLA-DR, CD3, CD19, CD56, CCR2 and CX3CR1. Antibody details are listed in the Supplemental Materials. Red blood cells were lysed using Pharm Lyse (BD Biosciences), followed by two wash steps using a BD Lyse wash assistant machine (BD Biosciences). Analysis was performed using a BD LSR Fortessa machine with FCS Express 6 (De Novo Software).

### Mouse Model of Myocardial Ischaemia and Reperfusion

All animal experiments were performed under EU legislation and approved by the animal ethics committee of Newcastle University. Acute myocardial infarction was induced in adult (12-14 weeks of age) male mice as previously described ^14^, except that hearts were reperfused after 60 minutes of ischaemia. Anaesthesia (3% isoflurane/97% oxygen) was maintained throughout the surgery and hypnorm (fluanisone/fentanyl) 0.2ml/Kg via subcutaneous injection was used as a pre-med. Breathing was maintained using mechanical ventilation via endotracheal intubation. Left-side thoracotomy was performed through the fourth intercostal space. The left anterior descending (LAD) coronary artery was ligated using a 7-0 prolene suture over a small piece of PE-10 tubing, approximately 1-2mm from the left atria. Myocardial ischemia was verified by blanching of the left ventricular myocardium. The chest wall was then temporarily closed and reopened after 60 minutes of ischaemia, followed by removal of PE tubing to allow reperfusion of the LAD artery. Sham-operated mice underwent thoracotomy and pericardial opening but no LAD artery ligation. After completion of the procedure, the chest was closed and mice were administered 0.1mg/kg buprenophrine subcutaneously and again at 24hours post-surgery to alleviate post-operative pain.

### Mouse Blood Sampling and Flow Cytometry Analysis

At the indicated times, murine blood (30μl) was collected from the tail vein into EDTA tubes (Teklab, #K100PP). Samples were incubated with Fc block (1/10 dilution) for 10 minutes and then with fluorescently labelled antibodies against CD11b, CD115, and Ly6C, for 30 minutes at room temperature in the dark. Antibody details are listed in the Supplemental Materials. Following red blood cell lysis (Pharm Lyse, BD) cells were resuspended in 300µl FACS buffer with the addition of 30µl DAPI (BD Pharmingen, #564907) immediately prior to data acquisition using an LSR Fortessa (BD). To acquire absolute cell counts, cellular events were calculated relative to bead events using TruCount beads (BD Biosciences, #340334).

### Tissue Processing and Immunofluorescent Staining

At the indicated times following reperfusion, mouse heart tissue was harvested and fixed in 1% paraformaldehyde prior to transfer into a 30% sucrose solution and embedding in Optimal Cutting Temperature compound on dry ice. Transverse 10um sections were collected on Superfrost-plus slides (VWR). Sections were blocked for 2hrs with 1% BSA, 5% donkey serum and 0.5% tween-20. Antibodies (anti-CD11b, anti-GFP and anti-podocalyxin) were incubated on sections overnight at 4°C. Following PBS washes, fluorescently-conjugated donkey secondary antibodies were added for 2 hours at room temperature in the dark. After PBS washes, coverslips were mounted using Prolong Gold antifade reagent with DAPI and left at 4°C overnight. Antibody details are listed in Supplemental Materials. Images were acquired using an M2 Axio Imager (Zeiss) and analysed using ZEN 2.3 software (Zeiss). The ischaemic region of the left ventricle was compared with the same region of sham hearts.

### Statistics

All statistical analysis was performed using SPSS and GraphPad Prism. Data were checked for Gaussian distribution using the Shapiro-Wilk test. Parametric tests were used to analyse normally distributed data and non-parametric tests for non-normally distributed data as indicated in the figure legends. Where multiple comparisons were made, reported values were corrected for multiple tests. Mean cell counts per patient group were used to calculate the average change over time. Data are presented as mean ± standard error of the mean except where otherwise stated. A p value less than 0.05 was considered significant.

## Results

### Non-classical monocytes are preferentially depleted in the circulation of STEMI patients at 90 minutes following reperfusion

A previous study originally designed to investigate lymphocyte responses by flow cytometry (Boag, Das et al. 2015) enabled retrospective interrogation of monocyte subset responses in 51 STEMI patients (Table 1) at the time of PPCI, and at 15, 30 and 90 minutes, and at 24 hours post-reperfusion. Analysis of the same monocyte populations in non-ST elevation myocardial infarction (NSTEMI) patients (n=15) enabled discrimination of acute cardiac ischaemia/reperfusion (I/R) injury effects from those caused by a more chronic ischaemic condition. Both patient groups underwent a similar clinical intervention using PPCI. Flow cytometry gating of monocyte subsets was based on forward and side scatter to define the monocyte gate, CD3 and CD56 negative staining to remove contaminating T-cells and NK-cells, and CD16 expression to define the different monocyte subpopulations (Figure 1A). STEMI patients showed a rapid fall of 45.4±4% (p<0.0001) in CD16^++^ NC monocytes from immediately prior to PPCI to 90 minutes post-reperfusion (Figure 1B). CD16^-^ classical and CD16^+^ intermediate monocytes showed a much smaller drop of 16.3±5% (p=0.023) and 23.8±5% (p=0.044), respectively, over the same time period (Figure 1B). By 24 hours post-reperfusion, all three monocyte populations show a clear increase in the circulation, likely due to the release of monocytes from the splenic and bone marrow reservoirs^12, 13^. To determine whether there was a specific effect of the PPCI procedure, a group of 15 NSTEMI patients was included as a control group. The numbers of classical and intermediate monocyte subsets are very similar between STEMI and NSTEMI patients (Figure 1B). However, there is a 25.8± 7% (p<0.05) fall in NC monocytes in NSTEMI patients, potentially as result of the PCI procedure, but the percentage fall is approximately half that seen in STEMI patients, suggesting there is additional NC monocyte depletion in response to cardiac ischaemia reperfusion injury. It is notable that NSTEMI patients have significantly increased counts of NC monocytes post-reperfusion compared with STEMI patients potentially as a result of their more prolonged (although milder) ischaemic status prior to PPCI. In a subset of STEMI patients (n=20), follow-up venous blood samples were collected at 3 to 6 months post-PCI, and revealed that the supra-baseline numbers of classical and intermediate monocytes seen at 24 hours had fallen back to baseline, whilst the NC monocytes (which had already returned to baseline numbers by 24h) remained at this level at 3-6 months (Supplementary Figure II).

**Figure 1.**
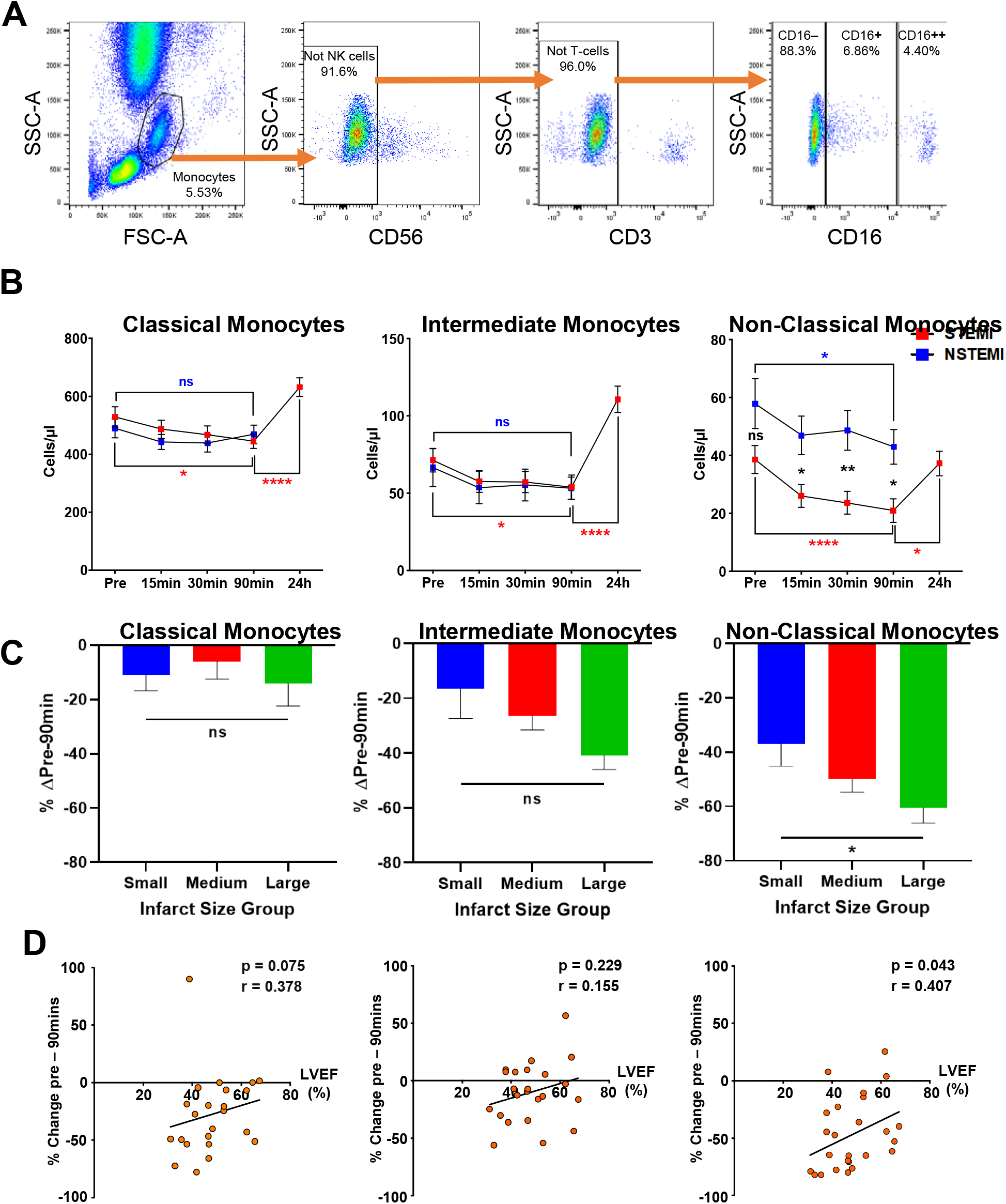
STEMI patient non-classical monocyte dynamics following PCI are associated with infarct size and LVEF. **A**. Flow Cytometry retrospective gating strategy for monocyte subset analysis in STEMI (n=51) and NSTEMI (n=15) patient blood. Quantification of monocytes was achieved using the scatter properties of monocytes, excluding T-cells (CD3+) and NK-cells (CD56+), followed by CD16 expression to define each monocyte subset (CD16-classical, CD16+ intermediate, CD16++ non-classical monocytes). **B**. Circulating monocytes (absolute counts) in STEMI and NSTEMI patients pre- and post-reperfusion. Black asterisks indicate statistical differences in counts between indicated time points in STEMI patients (data analysed by two-way mixed ANOVA with Bonferroni’s adjustment for multiple comparisons); Red and blue asterisks indicate statistical differences between counts at indicated time points in STEMI and NSTEMI patient groups, respectively. Data analysed by repeated measures paired analysis with Bonferroni adjustment for multiple comparisons as a full factorial model. * p<0.05, **** p<0.0001. **C**. Relationship between infarct size and total acute post-reperfusion period (pre-90min) in classical, intermediate, and NC monocytes in all STEMI patients that underwent cardiac MRI, categorized into tertiles based on infarct size (small: n=13, medium: n=16, large: n=13). Data analysed using one-way ANOVA with Tukey’s multiple comparison test. *p<0.05. **D**. LVEF significantly correlates with pre-reperfusion-90min changes in NC monocyte counts in STEMI patients with an anterior infarct (n=24). r=Spearman correlation coefficient.

### Acute post-reperfusion drop in circulating non-classical monocytes correlates with infarct size and cardiac function

We hypothesized that the transient depletion in NC monocytes at 90 minutes following reperfusion in STEMI patients reflects NC monocyte recruitment to the injured heart tissue and corresponds to the extent of cardiac injury. We therefore asked whether these dynamics correlated with infarct size (IS), MVO or myocardial function (measured by left ventricular ejection fraction, LVEF). These outcomes were assessed in the 42 patients who underwent cardiac MRI, which was between 1 and 8 days post-reperfusion. STEMI patients were first divided into three tertiles based on infarct size (Table S1 in Data Supplement shows group characteristics). Analysis of the dynamic changes in cell counts between each of these tertiles showed that STEMI patients with larger infarcts had a greater decline in NC monocytes between baseline (pre-reperfusion) and 90 minutes (post-reperfusion), compared with patients that had smaller infarcts (p=0.04) (Figure 1C). To examine correlation with cardiac function we limited the analysis to anterior STEMI patients (n=24) because left ventricular function is more severely compromised if the anterior rather than the inferior wall is involved ^15^. The fall in NC monocytes in anterior STEMI patients significantly correlated with LVEF (Spearman coefficient = 0.41, p=0.043; Figure 1D), potentially reflecting a specific response of NC monocytes to the level of cardiac injury. As ejection fraction generally falls over time, we verified that this correlation did not simply reflect time of MRI (data not shown). In contrast, dynamics of classical and intermediate monocytes showed no association with infarct size or LVEF. In addition, there was no association between dynamics of any monocyte subset and MVO (Supplementary Figure III), suggesting that monocyte populations do not have a contributory role in occlusion of the microvasculature in STEMI patients.

### Acute post-reperfusion fall in circulating non-classical monocytes in STEMI patients is confirmed in a second independent study

The flow analysis in the retrospective study was originally designed to examine lymphocyte dynamics, and monocytes were detected on the FSC/SSC gate, with any contaminating NK and T cells removed using CD56 and CD3, respectively (Figure 1A).. We therefore aimed to validate the monocyte subset dynamics using a more comprehensive panel of leukocyte markers in a prospective study. Taking a new STEMI patient cohort (n=13) (Table 1), we defined monocytes as CD14^+^ CD3^-^ CD19^-^ CD56^-^ and HLA-DR^+^. Monocytes were then further characterised as classical (CD14^++^ CD16^-^ CCR2^hi^ CX_3_CR1^lo^), intermediate (CD14^++^ CD16^+^ CCR2^mid^ CX_3_CR1^hi^) and NC (CD14^+^ CD16^++^ CCR2^lo^ CX_3_CR1^hi^) in line with a recent consensus document^16^ (Figure 2A). In this prospective analysis, we observed that classical, intermediate, and NC monocyte counts are similar between aortic, culprit, and non-culprit artery samples suggesting these pre-reperfusion measurements are not altered by local events close to the site of coronary occlusion (Supplementary Figure IV). Overall, the findings from this prospective study are largely in agreement with those from the retrospective study. Circulating NC monocyte counts significantly drop by 51.7± 7% (p=0.0008) from pre-reperfusion to 90 minutes. There is no further fall at 180 minutes suggesting that 90 minutes corresponds to the nadir of NC monocytes (Figure 2B). However, we noted that the absolute counts of NC monocytes are generally higher in the prospective study compared with the retrospective study. Further analysis revealed that many NC monocytes have lower side scatter than classical and intermediate monocytes, and overlap with the lymphocyte population on the forward/side scatter plot (Supplementary Figure V). Therefore these cells would have been excluded from flow cytometry analysis when using a simple FSC/SSC monocyte gating strategy due to their smaller size and lower granularity. Nevertheless, despite the different gating strategies used in the two studies, the percentage fall in NC monocytes between pre-and 90 minutes post reperfusion was strikingly similar (45.4±4% vs 51.7±7%). In STEMI patients in the prospective study, the mean number of classical and intermediate monocytes respectively show a 10.0±13% and 2.7±18% reduction in the blood from ischaemia to 90 minutes post-reperfusion (Figure 2B). This was a similar marginal fall to that seen in the retrospective study but was not statistically significant, likely due to the smaller size of the prospective patient cohort. Thus, we conclude from two independent studies that circulating NC monocytes in STEMI patients show a mean drop of approximately 50% at 90 minutes post-reperfusion. We also find that the extent of this fall correlates with the extent of cardiac injury (infarct size and LVEF) that may reflect the degree of monocyte margination to the coronary vessel endothelium of the infarcted heart tissue.

**Figure 2.**
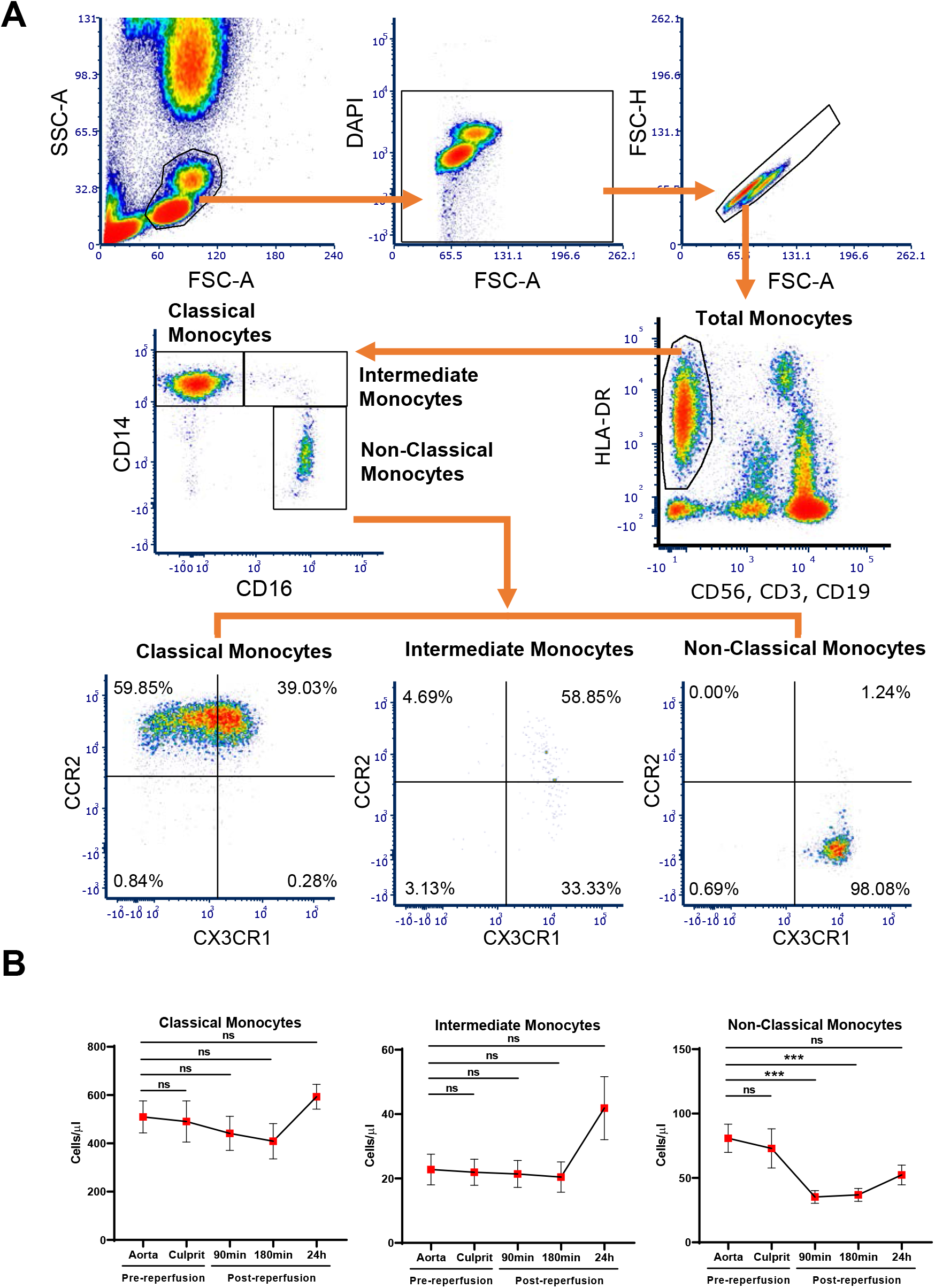
Prospective STEMI patient study confirms depletion of non-classical monocytes at 90 minutes post-reperfusion. **A**. Flow cytometry analysis using multi-parameter gating of blood samples from STEMI patients. Mononuclear cells based on forward (FSC-A) and side (SSC-A) scatter properties were characterised as live (DAPI-) single cells with T cells, B cells and NK cells excluded on the basis of CD3, CD19 and CD56 expression, respectively. HLA-DR+ CD14-cells were also excluded. Each HLA-DR+ monocyte subset was then defined as classical [CD14++ CD16-], intermediate [CD14++ CD16+] and NC [CD14+ CD16++]. Note that NC monocytes expressed higher levels of CX3CR1 and lower levels of CCR2 than classical monocytes, as expected. **B.** STEMI patients show similar circulating monocyte subset counts in blood from the aorta and culprit artery at baseline, but there was significant depletion of NC monocytes at 90 and 180 minutes post-reperfusion. Statistical analysis of blood samples at pre-reperfusion, 90min and 180min (all n=13), and 24h post-PCI (n=11) was performed using a mixed effect model with Dunnett’s adjustment for multiple comparisons. *** p<0.001.

### Monocytes marginate to the coronary vessels in the reperfused ischaemic murine heart

To track monocyte adhesion and extravasation in the heart and circulation immediately following myocardial I/R we used a mouse model of cardiac I/R injury. Acute myocardial infarction (MI) was created as previously described^14^, except that hearts were reperfused after 60 minutes of ischaemia. Sham-operated mice were used as controls. As STEMI patients showed a fall that was stable between 90 and 180 minutes post reperfusion and recovered by 24h, we monitored peripheral monocyte dynamics in the mouse at pre-reperfusion, and at 2h and 24h post-reperfusion. Circulating murine monocytes were quantified by flow cytometry as classical (CD11b+ CD115+ Ly6C^hi^), intermediate (CD11b+ CD115+ Ly6C^mid^) and non-classical (CD11b+ CD115+ Ly6C^lo^) monocytes (Supplementary Figure VI). Intermediate monocytes in mice have been characterised as having a heterogeneous transition phenotype between Ly6C+ and Ly6C-monocytes^17^. At 2h post-reperfusion, all circulating murine monocyte subsets showed a similar fall from pre-reperfusion levels in both sham operated and in cardiac I/R mice (Supplementary Figure VI). Therefore, there is a rapid monoctyopenia response to thoracotomy in the surgical mouse model, which cannot readily be separated from any specific response to cardiac I/R injury. In all cases (sham and cardiac I/R) circulating numbers of all three monocyte populations returned to pre-reperfusion levels by 24h, in line with known monocyte release from splenic and bone marrow stores in response to injury. Therefore, to investigate whether there was any specific monocyte recruitment to the injured myocardium following cardiac I/R injury we directly analysed heart tissue by immunofluorescent staining following cardiac I/R. We took advantage of heterozygous Cx3cr1-eGFP mice ^18^ to track monocytes in combination with the myeloid marker CD11b. In this mouse line the eGFP coding region replaces an essential part of one of the *Cx3cr1* alleles allowing it to serve as a reporter line for *Cx3cr1* expression, whilst *Cx3cr1* gene function is preserved in the corresponding wild-type allele. These mice express eGFP in all monocytes, but not in neutrophils ^18^. Thus monocytes in injured heart tissue were identified as CD11b+ GFP+ cells, neutrophils as CD11b+ GFP-cells and podocalyxin was used to identify endothelial cells. Adhesion of monocytes and neutrophils to coronary endothelial cells was observed in venous, but not arterial vessels, in line with the known role of the post-capillary venule as the primary site of leukocyte extravasation (Figure 3A). Furthermore, a significant increase in monocyte adhesion to the endothelium was seen in I/R hearts at 2 hours post reperfusion, compared with sham hearts (Figure 3A,D). This finding is consistent with a specific and rapid monocyte endothelial adhesion response to cardiac I/R injury. There were no vessel-adherent monocytes observed in naïve hearts (not shown). We noted that at 2h post reperfusion a small number of monocytes had already extravasated, but were still located at perivascular sites and had not yet migrated into the injured cardiac tissue, allowing them to be clearly discriminated from resident macrophages present in all hearts (Figure 3A). Although the numbers of marginated monocytes at 2h post reperfusion are lower than those of neutrophils, it is clear that both these cell populations respond very rapidly to injury (Figure 3A,D,E). By 24h post-I/R, there is a massive leukocyte infiltrate in the ischaemic region of the heart (Figure 3B). This infiltrate was dominated by neutrophils as expected, with monocytes a minority population (not shown), whereas by 3 days post-reperfusion, monocytes far outnumbered neutrophils (Figure 3C,F) in agreement with previous work^19^. Thus, murine monocytes rapidly marginalise to the coronary vessel endothelium within 2h of cardiac I/R injury and thereafter infiltrate into the heart tissue.

**Figure 3.**
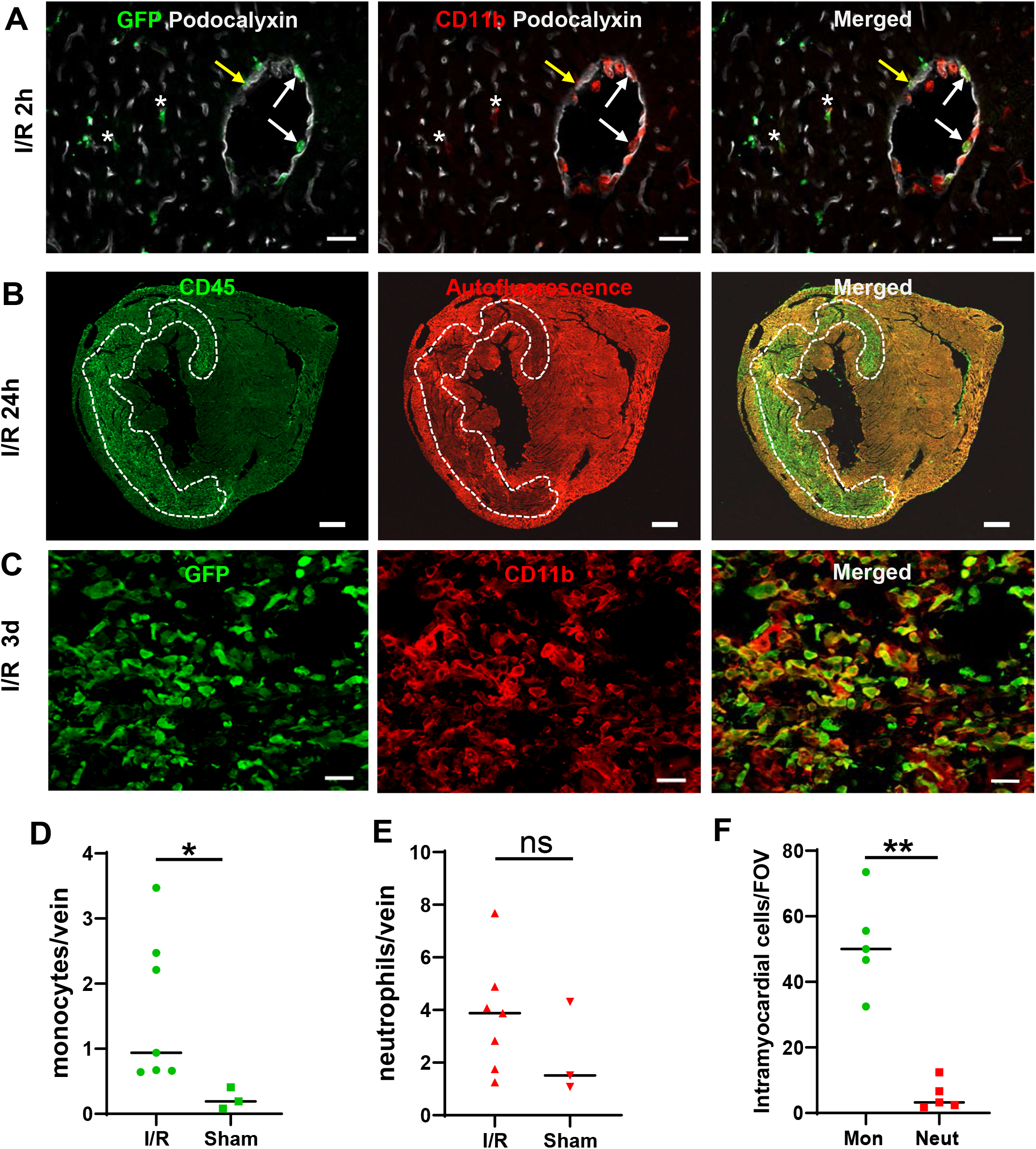
Murine monocytes show increased margination to the coronary vessel endothelium at 2 h post reperfusion. **A:** Monocytes (identified in the CX3CR1-GFP mouse as GFP+ CD11b+ cells, white arrows) and neutrophils (GFP-CD11b+ cells) intimately associate with venous endothelium (labelled with anti-podocalyxin) at 2h post reperfusion. Early extravasation of a monocyte can be seen (yellow arrow), and resident GFP+ CD11b+ macrophage are visible in the myocardium (asterisks). Scale bar = 20µm **B**: At 24h post reperfusion large numbers of CD45+ leukocytes locate to the infarcted region (dotted line), which is also characterised by loss of autofluorescence of the myocardium at extended exposure. Scale bar =500µm. **C:** At 3 days post reperfusion monocytes (GFP+ CD11b+) dominate the infarcted region. Scale bar = 20µm. **D-E:** Quantification of adherent monocytes and neutrophils in the injured heart at 2 hours post-reperfusion. Significantly higher numbers of monocytes adhere to the vascular endothelium of coronary vessels in the injured left ventricle of I/R hearts (n=7 hearts, 396 vessels), compared with sham controls (n=3 hearts, 186 vessels). In contrast there is no significant difference in adherent neutrophils between injured and sham hearts. Data analysed by Mann Whitney U test; *p<0.05. **F:** By 3 days post reperfusion there are high numbers of extravasated monocytes within the infarcted myocardium. Data analysed by Mann Whitney U test; **p<0.01.

## Discussion

We show the rapid depletion of circulating NC monocytes from pre-reperfusion to 90 minutes post-PCI in STEMI patients directly correlates with larger infarct size and impaired LVEF. Following MI, increased infarct size and decreased cardiac function are powerful predictors of later adverse cardiac outcomes including heart failure^20-22^. The ability to detect informative dynamic changes in monocytes immediately following reperfusion has potential clinical relevance since this time frame offers the potential to therapeutically intervene while STEMI patients are still hospitalized.

It is interesting to consider why the fall in NC monocytes (by 90 minutes post-reperfusion) is greater than that of classical and intermediate monocytes. Evidence from murine studies have shown that in steady state, NC monocytes actively engage the CX3CR1 receptor on their surface with FKN ligand on the endothelium to constitutively patrol the vasculature, acting on ‘stand-by’ for inflammatory stimuli^23-25^. The release of soluble fractalkine (sFKN) from the vascular endothelium by the metalloproteinase ADAM17 is triggered by inflammatory stimuli following cardiac I/R and leads to a cytokine gradient that would recruit CX3CR1 expressing monocytes to the injured tissue. Arrest and adhesion of NC monocytes to the vascular endothelium enables subsequent extravasation into the injured myocardial tissue. Consistent with this sequence of events, the nadir of NC monocytes at 90 minutes post-reperfusion coincides with the timing of the peak concentration of sFKN in the serum of STEMI patients ^3^. Thus, myocardial infarction likely drives increased NC monocyte recruitment from the peripheral blood to the injured site via a gradient of sFKN. Previous real time imaging in the mouse has shown that monocytes are recruited to the cardiac vasculature as early as 30min after permanent MI, although their specific location could not be discriminated ^26^. In contrast, we were able to show that murine monocytes adhere to the venous endothelium and initiate extravasation into the tissues by 2h post reperfusion. Unfortunately, we were unable to discriminate NC from classical monocytes in mouse heart tissue. We tested both CCR2 expression (high on classical monocytes) and PD-L1 expression (recently identified to be specific for NC monocytes^27^) to discriminate murine classical from NC monocytes, but found expression differences to be small and therefore insufficient to discriminate monocyte subsets in our hands. Furthermore, in contrast to human monocytes, CX3CR1 protein expression levels are similar across all murine monocyte subsets ^28^. Thus, any CX3CR1-dependent response would be similar in mouse monocytes irrespective of classical and NC status. The immediate monocyte responses in injured human I/R hearts are not known due to the very limited availability of tissues, but will likely follow a similar course.

A key question that our data raise is why a loss of NC monocytes from the circulation and recruitment to the coronary vessels in the affected heart tissue would correlate with decreased heart function. This is at first sight counter intuitive as classical monocytes are associated with an inflammatory phenotype whilst NC monocytes have a more reparative phenotype. However, there is evidence pointing to some pro-inflammatory and therefore potentially detrimental roles of NC monocytes. Stimulated NC monocytes express increased levels of the pro-inflammatory cytokine TNF-α compared with classical monocytes ^29^, and NC monocytes can also activate endothelial cells to recruit neutrophils ^30^. It is therefore possible that an early increased recruitment of NC monocytes to the injured myocardium following I/R may further exacerbate local inflammation and increase myocardial damage contributing to further deterioration of cardiac function. The rapid loss of NC monocytes from the circulation in STEMI patients immediately after successful reperfusion of the culprit coronary artery may therefore provide a rapid biomarker to prioritize patients for follow up, when there is increased opportunity to initiate effective interventions before patient hospital discharge.

Although previous studies showed monocytosis in MI patients in the days following PCI are associated with an increased risk of adverse outcomes ^7-11, 31^, to our knowledge, this is the first report of a relationship between circulating NC monocyte dynamics and myocardial injury within such a short time period (90 minutes) of PCI. As NC monocytes show the first detectable response amongst the monocyte subsets this finding is consistent with them being able to respond rapidly to tissue injury. In fact, CX3CR1 expression is high in human NC monocytes and enables a chemotactic response to FKN ligand, which is rapidly released at sites of injury and potentially explains their rapid loss from the circulation. On the other hand CCR2 expression is high on classical monocytes and enables these monocytes to respond to CCL2 ligand which accumulates more gradually at the injured site, consistent with a slightly later recruitment of classical monocytes to the injured heart at around day 3 where they play an important role in removing necrotic cells and cellular debris^32^.

Comparison of the two FACS protocols used for the two patient studies revealed that conservative gating of monocytes on the SSC/FSC plots to exclude leukocytes and neutrophils in the retrospective study meant that a proportion of the smaller NC monocytes within the lymphocyte SSC/FSC gate were missed (Supplementary Figure V). However, the detected NC monocytes in the retrospective study and the total NC monocytes in the prospective study showed a strikingly similar fall of 45% and 50%, respectively, from pre-reperfusion to 90 minutes post-reperfusion. We therefore consider the observed correlations between the acute drop of detected NC monocytes and heart function measurements in the retrospective study to be valid. In addition, there is a rapid margination of monocytes to the coronary vessel endothelium in a mouse surgical model of myocardial infarction, and although there are many similarities between mouse and human monocytes ^8 33^, there are also clear differences that may limit a direct translation of findings from mouse to human. For example, Lys6C is highly expressed in murine classical monocytes, but is not found in human monocytes. Furthermore, NC monocytes are a minor subpopulation (∼10%) of human monocytes, but a major subset of mouse monocytes, depending on background strain^34^. Thus, there is some caution needed when extrapolating findings found in mouse to the conditions in human. Nevertheless, we observed rapid monocyte margination to the coronary endothelium in the mouse heart at 2 hours post reperfusion potentially explaining the concomitant fall in circulating monocytes in STEMI patients. As this response is so rapid it is likely to precede synthesis of new cytokine proteins such as CCL2, it is therefore tempting to speculate that sFKN is rapidly released to recruit CX3CR1 expressing monocytes to the injured site.

In conclusion, STEMI patient non-classical monocyte dynamics immediately following PCI are associated with infarct size and left ventricular function, suggesting a mechanistic link between these cells and cardiac injury. Mouse monocyte adhesion to myocardial venous endothelium occurs over a similar time frame following reperfusion in a pre-clinical mouse model of transient cardiac ischaemia. Non-classical monocytes may therefore have an earlier role in the acute post-reperfusion period than previously realised, and offer a new rapid biomarker of cardiac I/R injury.

## Contribution

HMA, IS and SA designed and supervised the study; SAM, CP, RER, ES, LD, and SEB performed experiments and data analysis; SEB and LS collected clinical samples; HMA, CP and SM wrote the manuscript and all authors checked, edited and approved the manuscript for publication. We acknowledge the Newcastle University Flow Cytometry Core Facility (FCCF) for assistance with the generation of Flow Cytometry data.

## Abbreviations

ACS: acute coronary syndrome
CX3CR1: C-X3-C Motif Chemokine Receptor 1
DAMPs: damage-associated molecular patterns
FKN: Fractalkine ligand, also known as CX3CL1
LAD: left anterior descending
LVEF: left ventricular ejection fraction
MACE: major adverse cardiac events
MI: myocardial infarction
MVO: microvascular occlusion
NC: non classical
NSTEMI: non-ST elevation myocardial infarction
PPCI: primary percutaneous coronary intervention
ROS: reactive oxygen species
sFKN: Soluble fractalkine
STEMI: ST elevation myocardial infarction

