## Supplementary Materials for "Rapid fall in circulating non-classical monocytes in ST elevation myocardial infarction patients correlates with cardiac injury"

### Supplementary Figure I

A

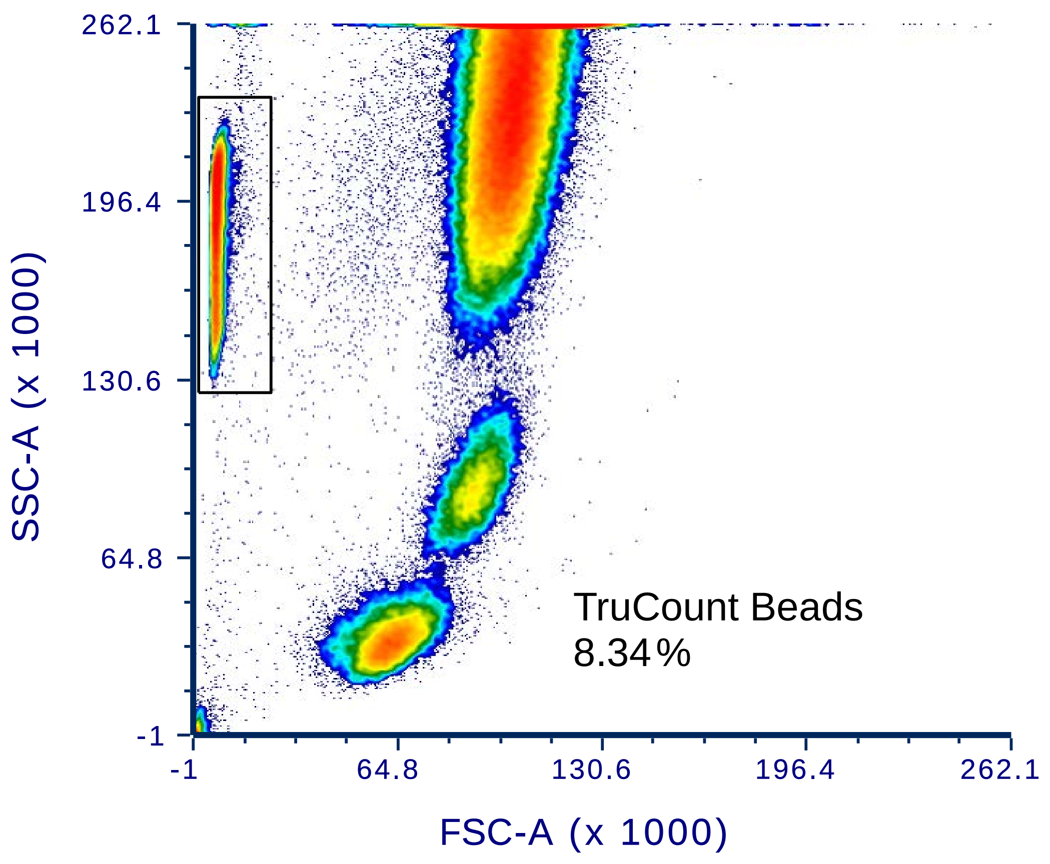


B

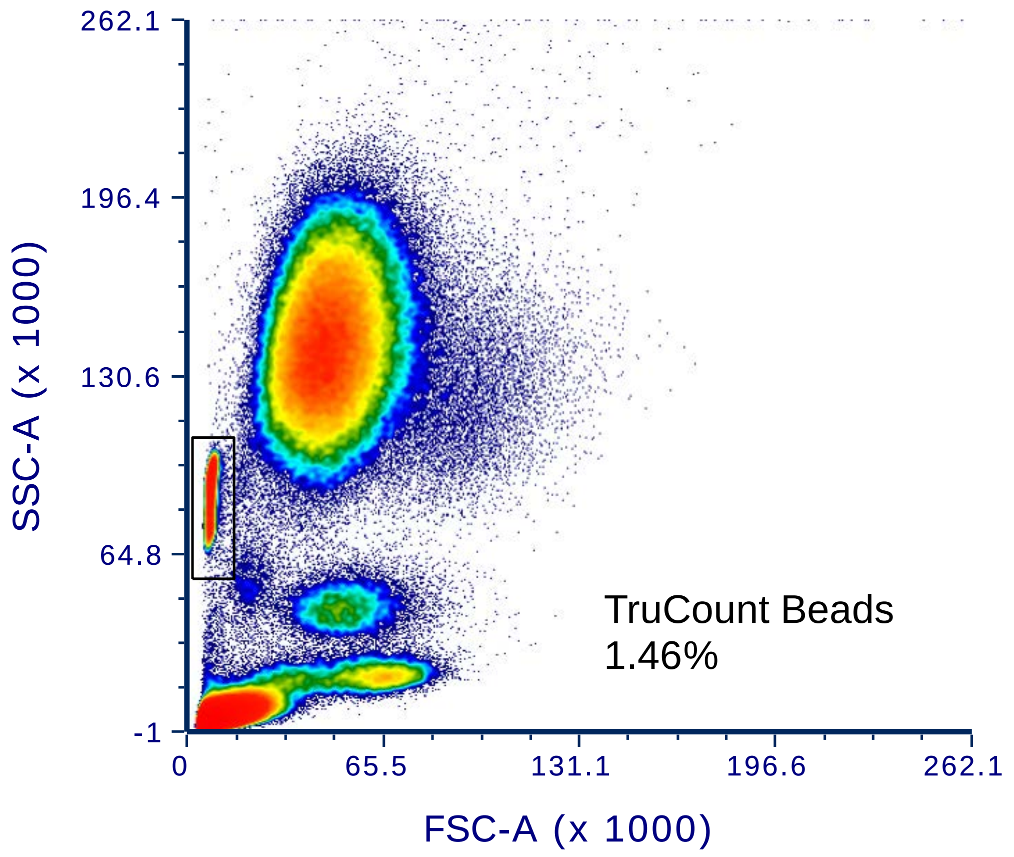


**Figure I. Flow cytometry plots to illustrate how Trucount beads are used to quantify STEMI patient leukocyte cell counts in retrospective study (A) and prospective study (B).**

### Supplementary Figure II

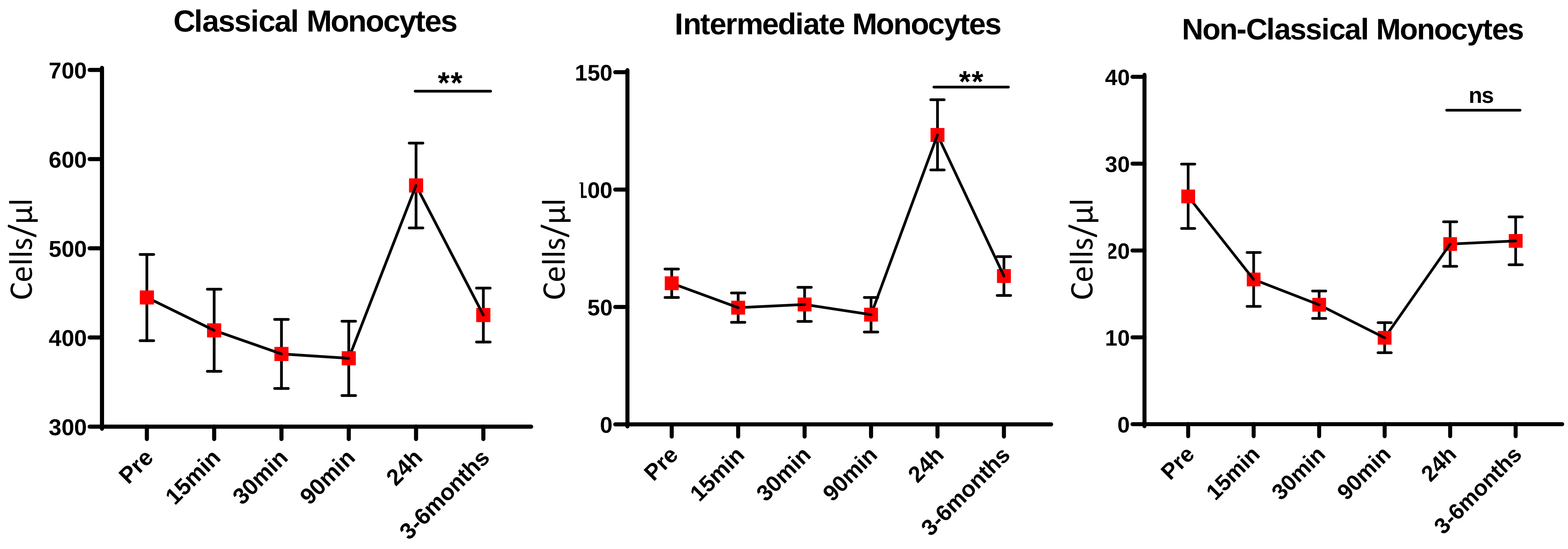


**Figure II. STEMI patient circulating monocyte counts at 3 month follow-up.** In a subset of STEMI patients (n=20), peripheral blood samples were collected at 3-6 months post-reperfusion. By this time point, $CD16^-$ classical, and $CD16^+$ intermediate monocyte counts returned from elevated levels seen at 24h post-reperfusion back to baseline levels. Non-classical monocytes remained at approximately baseline levels. Data are shown as mean +/- SEM and were analysed by repeated measures one-way Anova with Tukey's adjustment for multiple comparisons. ***p<0.001.

### Supplementary Figure III

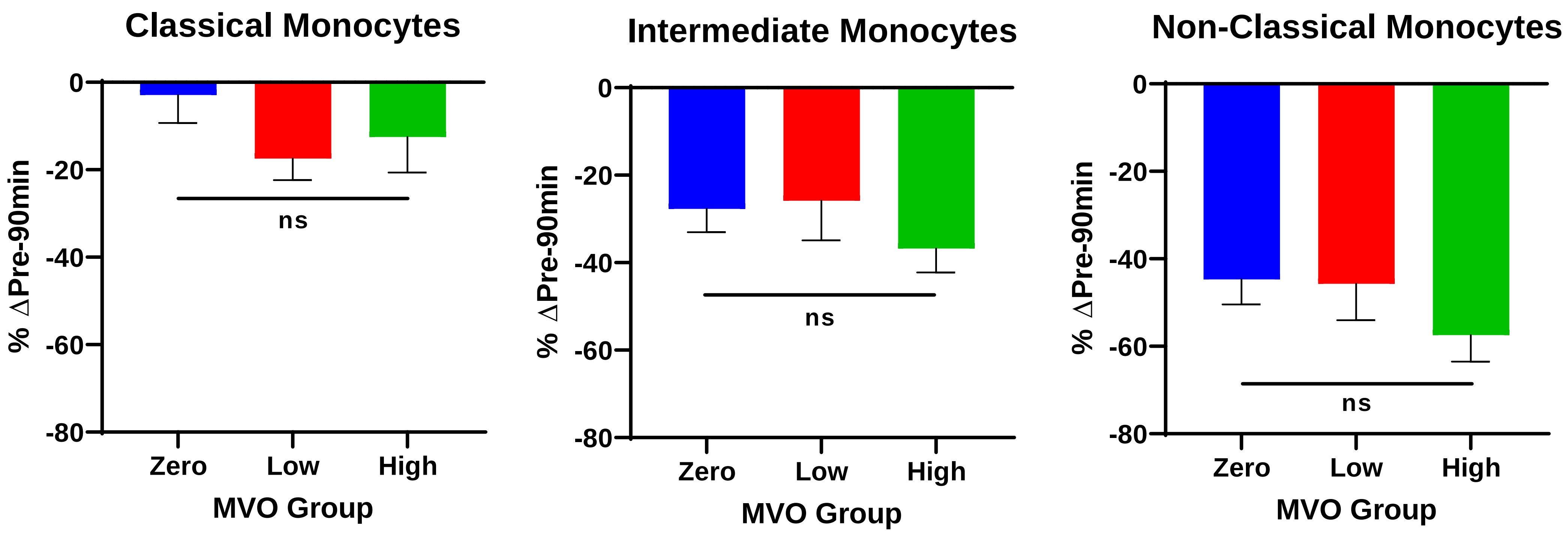


**Figure III. Relationship between MVO and total acute post-reperfusion changes (pre-90min) in classical, intermediate, and non-classical monocytes in STEMI patients.** STEMI patients underwent cardiac MRI to detect and quantify MVO. Data analysed by one-way ANOVA with Tukey's multiple comparison test. Patients were divided into tertiles based on mass of MVO as follows: none: n=17, low (0.1-2.7g): n=12, and high (>2.7g): n=13.

### Supplementary Figure IV

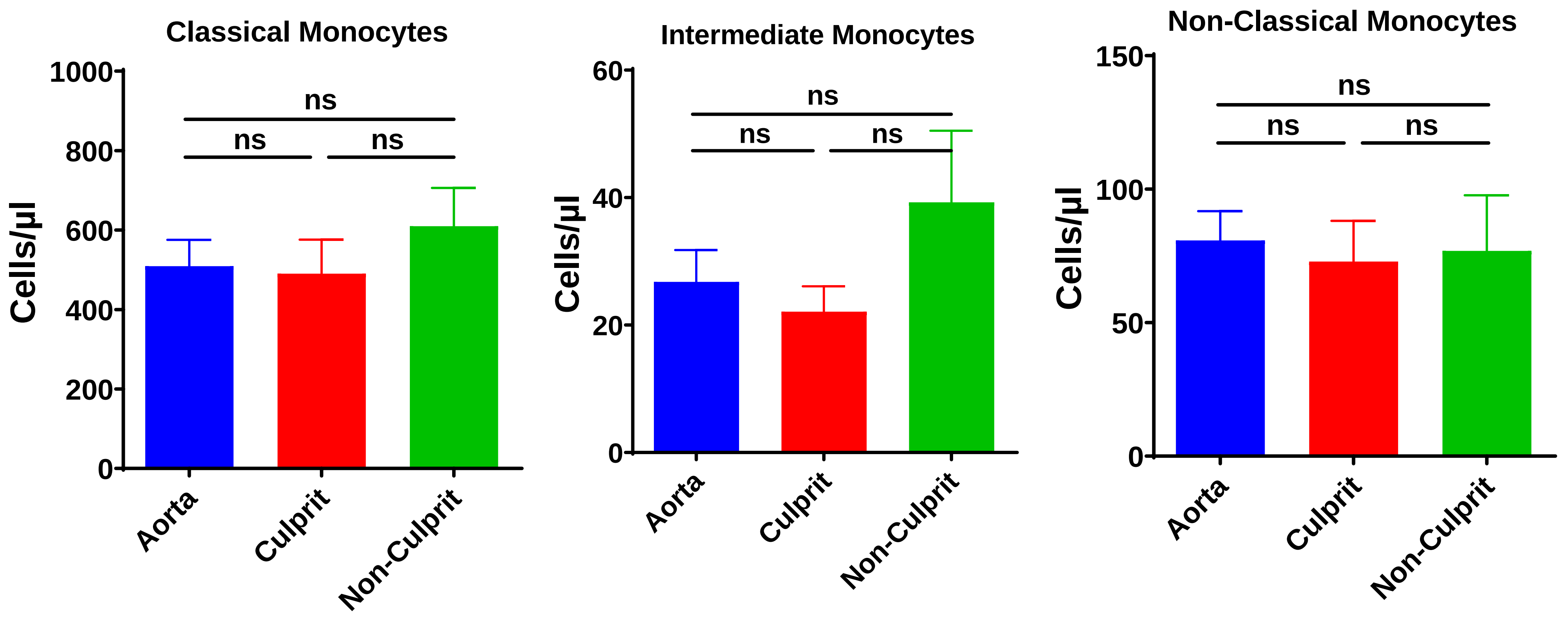


**Figure IV. STEMI patient monocyte subset counts taken immediately prior to reperfusion from different locations are similar.** For each STEMI patient, RCA and LCA were determined as the culprit or non-culprit artery. No significant differences in any monocyte subset counts were present between locations at the time of pre-perfusion. Data analysed by one-way ANOVA with Bonferroni's multiple comparison test. Aorta n=13, Culprit artery n=13, Non-Culprit artery n=5.

### Supplementary Figure V

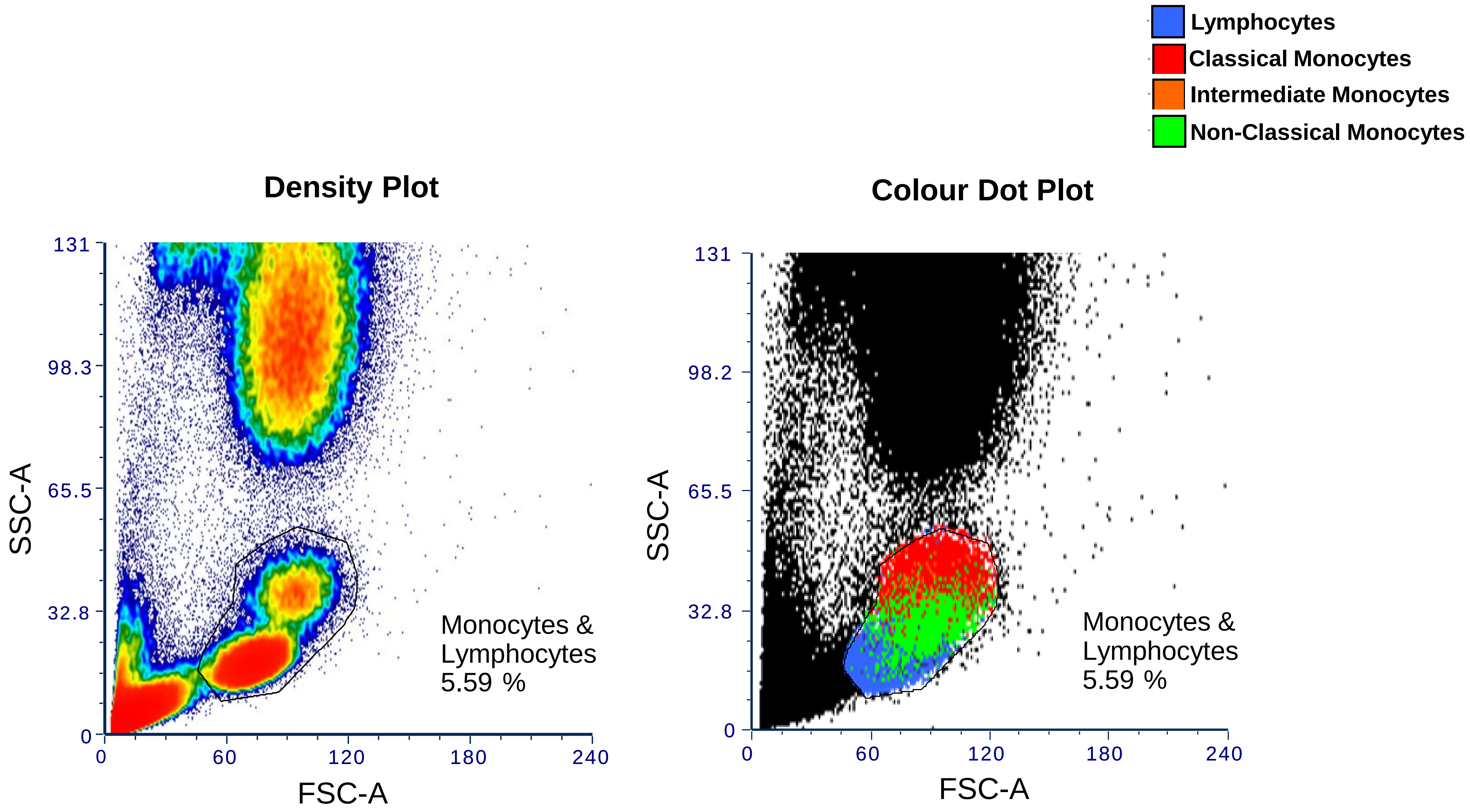


**Figure V. Comparison of gating strategies between retrospective and prospective studies.**
Using a colour dot plot to locate NC monocytes from STEMI patients analysed by the detailed gating method used in the prospective study reveals that NC monocytes (green) overlay with the lymphocytes on the SSC-A/FSC-A density plot.

### Supplementary Figure VI

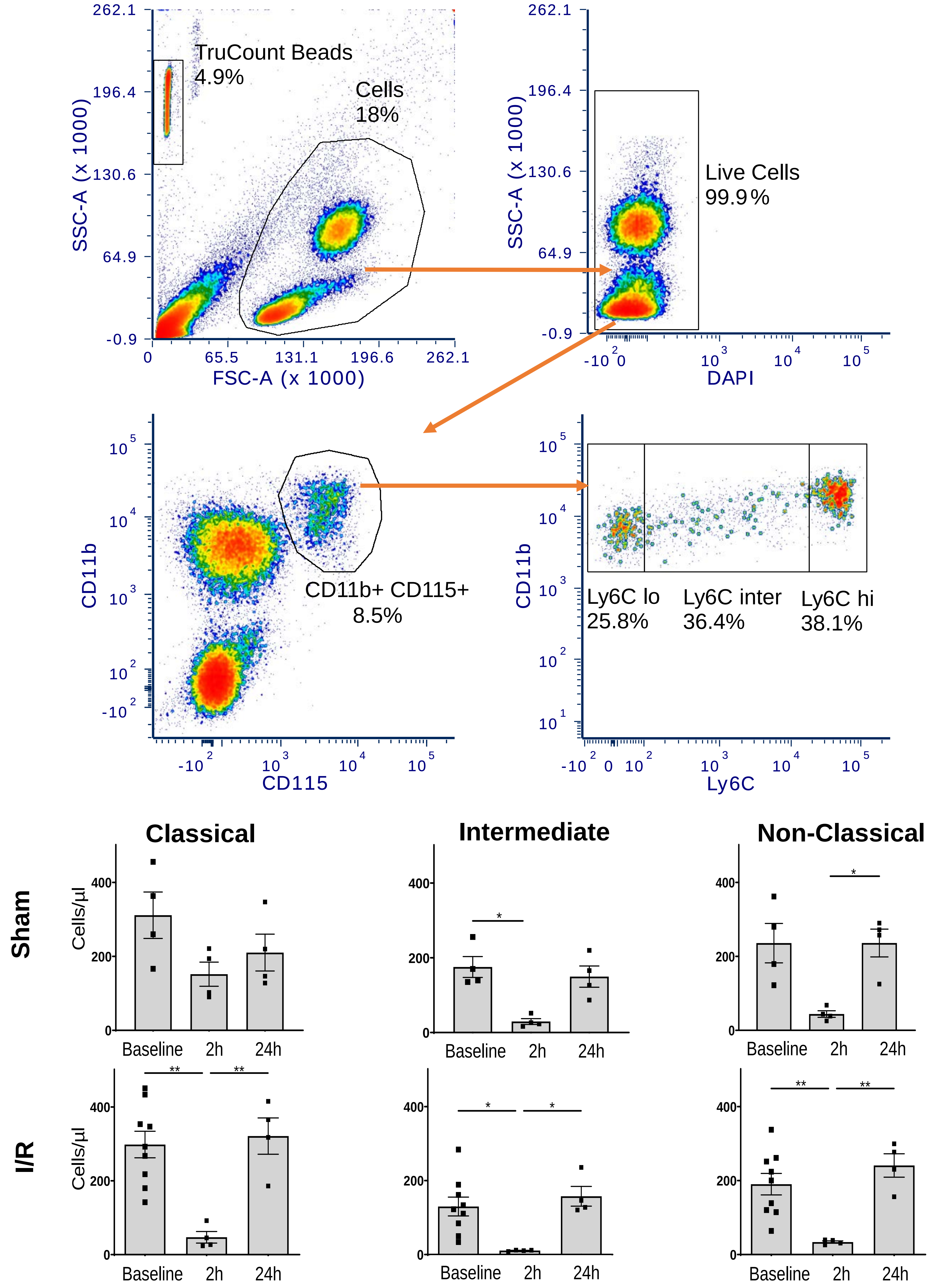


**Figure VI Circulating murine monocytes show a transient fall at 2 h post reperfusion in response to surgical thoracotomy**

A. Flow cytometry gating strategy to define mouse blood monocytes. Mouse circulating monocytes defined as classical (CD11b+ CD115+ Ly6Chi) intermediate (CD11b+ CD115+ Ly6Cmid) and non-classical (CD11b+ CD115+ Ly6Clo). B. Numbers of circulating monocytes before after surgery in sham and cardiac I/R mice. Circulating classical, intermediate and NC monocytes show a similar fall at 2h post-thoracotomy (sham, n=4/group) or 2h post-reperfusion (cardiac I/R n≥4/group) and then return to baseline levels by 24h. Data are analysed by one way ANOVA with Tukey post hoc test, * $p<0.05$; ** $p<0.01$.

### Table S1

| Infarct Size Group | | | | |
|---|---|---|---|---|
| | **Small (< 13.3 %)** | **Medium (13.3% - 23.8%)** | **Large (>23.8%)** | **p-value** |
| | **n = 13** | **n = 16** | **n = 13** | |
| **Patient characteristics** | | | | |
| **Age** | 13 | 16 | 13 | NA |
| **Sex (male:female)** | 57.6 ± 2.6 | 59.13 ± 2.3 | 57.9 ± 3.0 | 0.8927 |
| **BMI** | 10:3 | 13:3 | 12:1 | 0.5520 |
| **Diabetes mellitus** | 27.9 ± 1.4 | 27.1 ± 1.4 | 25.6 ± 1.0 | 0.4227 |
| **Family history of CAD** | 0 (0) | 2 (12.5) | 1 (7.7) | 0.4278 |
| **Active smoker** | 6 (46.2) | 7 (43.8) | 5 (38.5) | 0.9021 |
| **Hypertension** | 9 (69.2) | 8 (50.0) | 7 (53.8) | 0.5580 |
| **Anterior MI** | 5 (38.5) | 3 (18.8) | 2 (15.4) | 0.3210 |
| **Procedural characteristics** | | | | |
| **Door-to-balloon time (minutes)** | 26.5 ± 4.2 | 30.4 ± 4.8 | 24.0 ± 2.6 | 0.6481 |
| **Onset-to-reperfusion time (minutes)** | 168.2 ± 23.2 | 181.3 ± 27.2 | 136.6 ± 20.3 | 0.1837 |
| **MRI characteristics** | | | | |
| **Time to MRI (days)** | 2.6 ± 0.5 | 2.5 ± 0.4 | 3.0 ± 0.5 | 0.3923 |
| **LVEF (%)** | 57.8 ± 2.8 | 55.7 ± 2.0 | 44.8 ± 3.0 | 0.0065 ** |

**Table S1.** Relationship between infarct size and baseline data for STEMI patients that underwent cardiac MRI in the retrospective study. Continuous variables expressed as mean ± SEM; categorical variables expressed as n (%). Statistical analyses were performed using unpaired, two-tailed t-tests for continuous data, and Chi-squared tests for categorical data.

### Major Resources Table

#### Animals (in vivo studies)

| Species | Vendor or Source | Background Strain | Sex | Persistent ID / URL |
|---|---|---|---|---|
| Mouse (wild type) | Charles River | C57BL/6 | M | |

#### Genetically Modified Animals

| Species | Vendor or Source | Background Strain | Other Information | Persistent ID / URL |
|---|---|---|---|---|
| Mouse | Jax labs | C57BL/6 | $Cx3cr1^{tm1Lit}$ | http://www.informatics.jax.org/allele/MGI:2670351 |

#### Antibodies – Mouse Immunofluorescent Staining

| Target antigen | Vendor or Source | Catalog # | Working concentration | Persistent ID / URL |
|---|---|---|---|---|
| Rat Anti-Mouse CD11b | BD Pharminogen | 553308 | 1/100 | https://www.bdbiosciences.com/us/applications/research/stem-cell-research/mesenchymal-stem-cell-markers-bone-marrow/mouse/negative-markers/purified-rat-anti-cd11b-m170/p/553308 |
| Chicken Anti-Mouse GFP | Abcam | ab13970 | 1/300 | https://www.abcam.com/gfp-antibody-ab13970.html |
| Goat Anti-Mouse Podocalyxin (0.2mg/ml) | R&D systems | AF1556 | 1/100 | https://www.rndsystems.com/products/mouse-podocalyxin-antibody_af1556 |

#### Other – Mouse Immunofluorescent Staining

| Description | Source / Repository | Persistent ID / URL |
|---|---|---|
| ProLong™ Gold Antifade Mountant with DAPI [Catalog number: P36931] | ThermoFisher Scientific | https://www.thermofisher.com/order/catalog/product/P36931#/P36931 |

#### Antibodies – Mouse Blood FACS

| Target antigen | Vendor or Source | Catalog # | Working concentration | Persistent ID / URL |
|---|---|---|---|---|
| Anti-Mouse CD11b APC | BioLegend | 101211 | 1/50 | https://www.biolegend.com/en-gb/products/apc-anti-mouse-human-cd11b-antibody-345 |
| Anti-Mouse CD115 BV711 | BioLegend | AFS98 | 1/25 | https://www.biolegend.com/en-ie/products/brilliant-violet-711-anti-mouse-cd115-csf-1r-antibody-9030 |
| Anti-Mouse Ly6C PE | BioLegend | 128007 | 1/50 | https://www.biolegend.com/en-us/products/pe-anti-mouse-ly-6c-antibody-4904 |

DOI [to be added]

**Antibodies – STEMI Patient Blood FACS (Prospective Study)**

| Target antigen | Vendor or Source | Catalog # | Working concentration | Persistent ID / URL |
|---|---|---|---|---|
| Anti-Human HLA-DR BV421 | BioLegend | 307635 | 1/10 | https://www.biolegend.com/en-gb/search-results/brilliant-violet-421-anti-human-hla-dr-antibody-7226 |
| Anti-Human CD3 FITC | BioLegend | 300406 | 1/10 | https://www.biolegend.com/en-gb/products/fitc-anti-human-cd3-antibody-863 |
| Anti-Human CD19 FITC | BioLegend | 302206 | 1/10 | https://www.biolegend.com/en-gb/products/fitc-anti-human-cd19-antibody-717 |
| Anti-Human CD56 FITC | BioLegend | 318304 | 1/10 | https://www.biolegend.com/en-gb/products/fitc-anti-human-cd56-ncam-antibody-3795 |
| Anti-Human CD16 APC-H7 | BD Biosciences | 560715 | 1/10 | https://www.bdbiosciences.com/us/reagents/research/antibodies-buffers/immunology-reagents/anti-human-antibodies/cell-surface-antigens/apc-h7-mouse-anti-human-cd16-3g8/p/560715 |
| Anti-Human CD14 BV510 | BioLegend | 301842 | 1/10 | https://www.biolegend.com/en-gb/products/brilliant-violet-510-anti-human-cd14-antibody-8001 |
| Anti-Human CX3CR1 APC | BioLegend | 341609 | 1/10 | https://www.biolegend.com/en-gb/products/apc-anti-human-cx3cr1-antibody-6605 |
| Anti-Human CCR2 PE-Cy7 | BioLegend | 357212 | 1/10 | https://www.biolegend.com/en-gb/products/pe-cyanine7-anti-human-cd192-ccr2-antibody-8938 |

DOI [to be added]
